# Shared Genetic and Regulatory Mechanisms in Acute Myeloid Leukaemia and Blast Crisis Chronic Myeloid Leukaemia

**DOI:** 10.1101/2024.10.26.620392

**Authors:** Sulagna Basu, Rakesh Pandey

**Affiliations:** Quantitative and Systems Biology Lab, Bioinformatics, MMV, Banaras Hindu Univeristy, Varanasi-221005, India

## Abstract

Acute Myeloid Leukemia (AML) and Blast Crisis Chronic Myeloid Leukemia (BC CML) share several clinical characteristics, including the accumulation of immature myeloid cells and disrupted hematopoiesis. Despite the similarities, the two diseases exhibit different genetic profiles, contributing to distinct disease progressions and therapeutic outcomes. In this study, we perform a comparative analysis using bioinformatics and systems biology approaches to explore shared and unique mutational signatures, transcriptional networks, and molecular regulators in AML and BC CML. Our analysis identifies 29 genes common to both diseases, which are involved in key biological processes such as cell proliferation, cell death regulation, and transcriptional control. A subsequent pathway enrichment highlights common pathways, including the PI3K-AKT signaling and cytokine-mediated signaling, that may contribute to disease progression and resistance to therapy. Additionally, we construct a protein-protein interaction network and a transcription factor-miRNA coregulatory network, revealing key hub genes such as TP53, RUNX1, and BCL2. These genes could be potential shared therapeutic targets for treating AML and BC CML. Therefore, our analysis offers insights into the molecular underpinnings of these aggressive hematological malignancies.

## Introduction

Cancer is one of the major challenges of 21^*st*^ Century and according to World Health Organization it accounts to nearly 10 million deaths world-wide annually.^1^ Amongst them, Leukemia has prevalence of 2.5 percent across the globe.^2^ Leukemia is considered to be a cancer of blood progenitor cells that originate primarily in the bone marrow and the lymphatic system. In Leukemia, the abnormal production of immature or dysfunctional white blood cells as well as uncontrolled proliferation of all types of blood cells lead to crowding out normal blood cells and interfering with the body’s ability to fight infections, carry oxygen, and control bleeding^3–5^. Myeloid leukemias are a subtype of leukemia that affect the hematopoetic stem cells of myeloid lineage, which mature into red blood cells, platelets, and various types of white blood cells (except for lymphocytes). It is further categorized into Acute Myeloid Leukaemia (AML) and Chronic Myeloid Leukaemia (CML) based on how fast it progresses.

Despite having a similar nomenclature and common involvement of myeloid cells, both diseases do not exhibit similar pathogenesis^6–8^. AML involves a complex interplay of genetic and environmental factors leading to the uncontrolled proliferation of the myeloblast cells with consequence of granulocytopenia, anaemia, thrombocytopenia and leukostasis^9^. Relatively, CML has a simpler genetic factor - chromosomal translocation of chromosomes 9 and 22, known as the Philadelphia chromosome. In the chronic phase of CML, this genetic mutation leads to abnormal proliferation of the myeloid stem cells that still retain the capacity to differentiate into mature blood cells to a certain extent.^10,11^ Subsequent accumulation of other genetic mutations are considered to be responsible for the loss of the differentiation capacity of the leukemic stem cells defined as the final phase of CML, Blast Crisis phase^12,13^. In this phase, a rapid increase in number of Blast cells occur in both bone marrow and peripheral blood, which is a characteristics similar to that of AML^14^.

In CML, additional genetic and molecular abnormalities other than the original BCR-ABL translocation have also reported similar to the diverse mutations found in AML . Interestingly, sometimes BCR:ABL mutation could be found alongside other driver mutations in AML patients leading to adverse risk stratification, i.e. having a poor prognosis^15^. BC CML has a much lower median survival than that of AML. Given the poor prognosis and abysmal survival rate of BC CML, AML patients may have a better outlook depending on specific factors such as age, genetic disposition, mutational burden etc.^16^. Despite the observations of genetic overlaps between these two hematologic malignancies, contributing to their aggressive nature and similar clinical presentation, a comprehensive study to compare them is still missing. Here, we use a systems biology approach to compare and explore the mutational signatures of AML and Blast crisis CML. Such comparisons have been performed but in context to other diseases^17^. Our analysis suggests that the mutational signatures in BC CML resemble those observed in *de novo* AML, suggesting shared mutational processes driving progression to advanced phase of both the diseases. In addition, our study enlist shared transcriptional as well as miRNA regulations in both the diseases. Resulting networks suggest that regulation of disease associated genes for both the diseases have shared commonality and hence can share same therapeutic targets.

## Materials and Methods

### Generating gene lists common for AML and BC CML

Top 500 genes related to AML and BC CML were collected from CTD^18^, GeneCards^19^, and DisGeNET^20^ databases. All retrieved data were included if the raw data set had fewer than 500 entries. Duplicates were removed and lists were merged for genes involved for each disease from the three databases. Finally common genes were found to be shared amongst the two disease using union and intersect operators in Excel.

### Pathway enrichment analaysis

Using Enrichr^21^, a comprehensive gene set enrichment web tool, a series of enrichment analyses were carried out to obtain comprehensive information on describing cellular processes and pathways for signaling. Gene ontology (GO)^22^ analysis was performed on the list of common genes. To identify metabolic routes, the Kyoto Encyclopedia of Genes and Genomes (KEGG) pathway was utilized. In addition to the KEGG pathway^23^, WikiPathways^24^, Reactome^25^, and BioCarta^26^ databases were utilized to provide a more thorough understanding of the pertinent signaling pathways.

### Mutation analysis

We used Cancer Gene Census(CGC) from the COSMIC database^27^ to perfrom mutation analysis of the 29 common genes. CGC currntly has mutation data for 15 out of the 29 genes. This data includes whether there are known somatic and germline mutation of the genes in relation to tumour types, the role it plays in cancer as well mutation type.

### Network construction and interaction between common gene list

Using the 29 common genes, STRING database^28^ was used to build a PPI network with all other settings set to default, and a minimum needed interaction score of 0.4 for the confidence score. Cytoscape (V3.8.2)^29^ was used to analyze and illustrate the PPI results. The Cytoscape plug-in CytoHubba^30^ was used to calculate degree topology and maximum clique centrality of the network and identify top ten hub proteins with the highest degree values.

### TF and miRNA mediated network of common genes

In order to examine the relationship between the common genes and miRNAs, as well as the effect of TFs on the expression and functional pathways of the common genes, different programs under NetworkAnalyst 3.0^31^ was utilized. The TRRUST database provided curated human transcription factor and gene target information^32^ for construction of transcriptional regulatory network. The miRTarBase (v9.0) database provided comprehensive experimentally validated miRNA-gene interaction data for constructing Gene-miRNA interaction network.^33^

### Putative Coregulation in AML and BC CML

The shared genes were submitted to NetworkAnalyst 3.0 again to construct a TF–miRNA coregulatory network. RegNetwork provided the information on regulatory interactions that had been curated based on the literature^34^. Due to excessive number of nodes, minimum connected network was selected to define the core network topology. Cytoscape was also used to visualize pertinent results.

## Results

### List of Genes common for AML and BC CML

We have enlisted the genes associated with AML and BC CML from publicly available databases: CTD, GeneCards, and DisGeNET. A summary of the list is shown in Table 1. After combining data from all the three databases and removing duplicates, we have found 251 genes linked to AML. With similar approach, we have found 157 genes linked to BC CML. We observed that there are 29 genes that are linked with both AML and BC CML (Fig. 1).

**Table 1.**
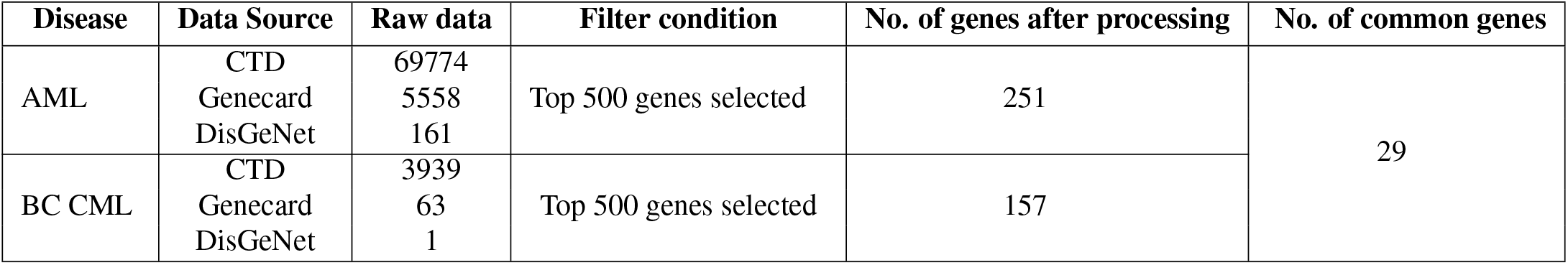
A summary of the list of genes associated with BC CML and AML from different Databases.

| Disease | Data Source | Raw data | Filter condition | No. of genes after processing | No. of common genes |
| --- | --- | --- | --- | --- | --- |
| AML | CTD | 69774 | Top 500 genes selected | 251 | 29 |
|  | Genecard | 5558 |  |  |  |
|  | DisGeNet | 161 |  |  |  |
| BC CML | CTD | 3939 | Top 500 genes selected | 157 |  |
|  | Genecard | 63 |  |  |  |
|  | DisGeNet | 1 |  |  |  |

**Figure 1.**
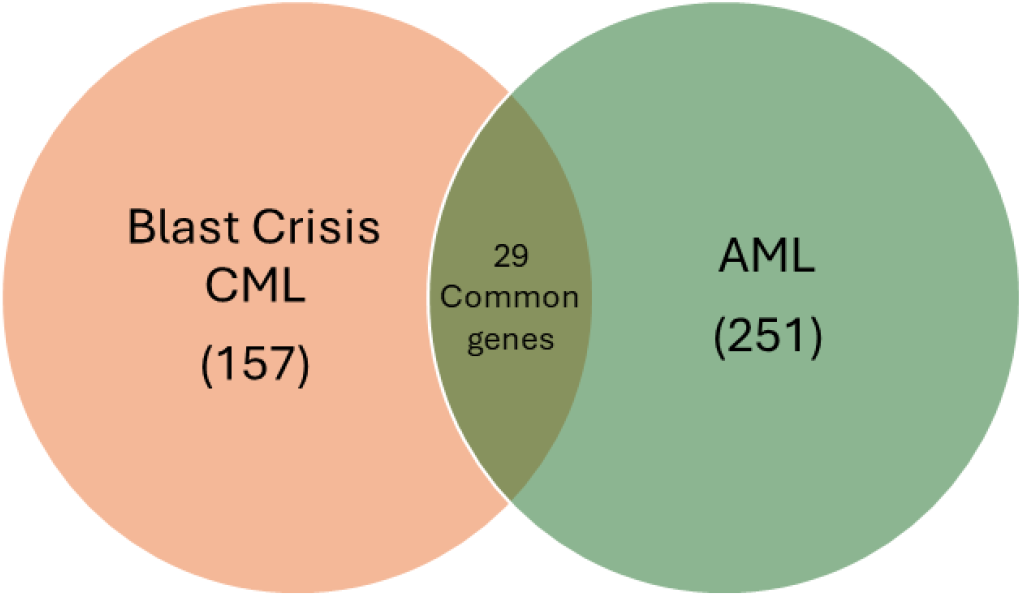
A venn diagram representing number of genes associated with BC CML and AML along with the numbers of genes common for the both diseases.

### Pathway enrichment analysis

Using the list of common 29 genes we have performed pathway enrichment analysis using a web based tool Enrichr that carried out the route of enrichment analysis, gene ontology, and visualized the result. The Enrichr calculates a composite score, that is based on the log of the Z-score and p-value. We have used GO database as an annotation source. The obtained gene ontology analysis of the 29 common genes could be categories in three groups: biological processes, cellular components, and molecular function. Following that we obtained the top pathways our common genes are involved in and are associtaed to with the diseases. (Fig. 2)

**Figure 2.**
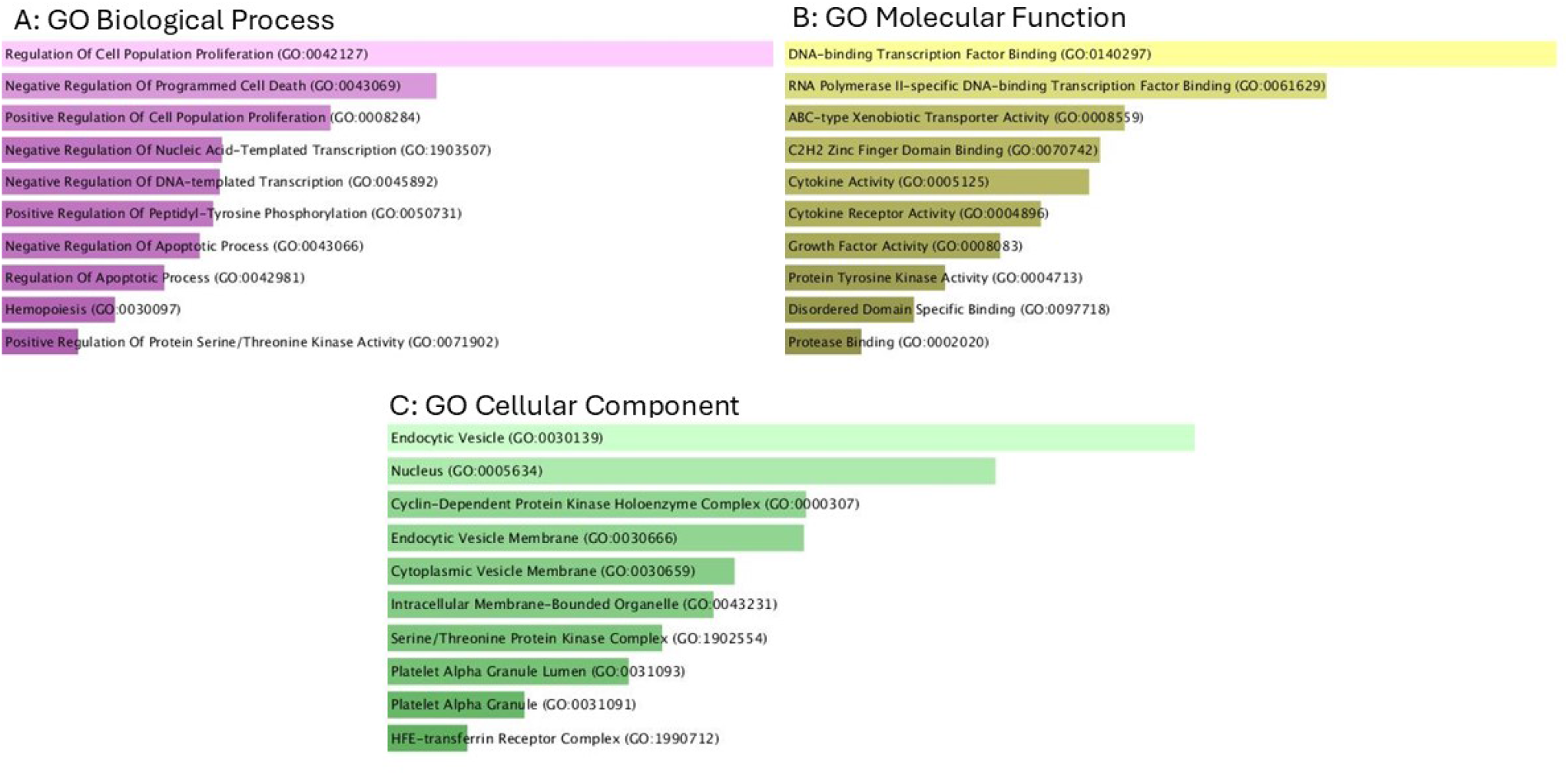
Gene Ontology terms for the 29 common genes under three categories: (A) Biological Processes, (B) Molecular Function, (C) Cellular Component.

Further, we obtained the most affected pathways of the common genes using four databases: KEGG^23^, WikiPathways^24^, Reactome^25^ and BioCarta^26^. Respective results are shown in (Fig. 3).

**Figure 3.**
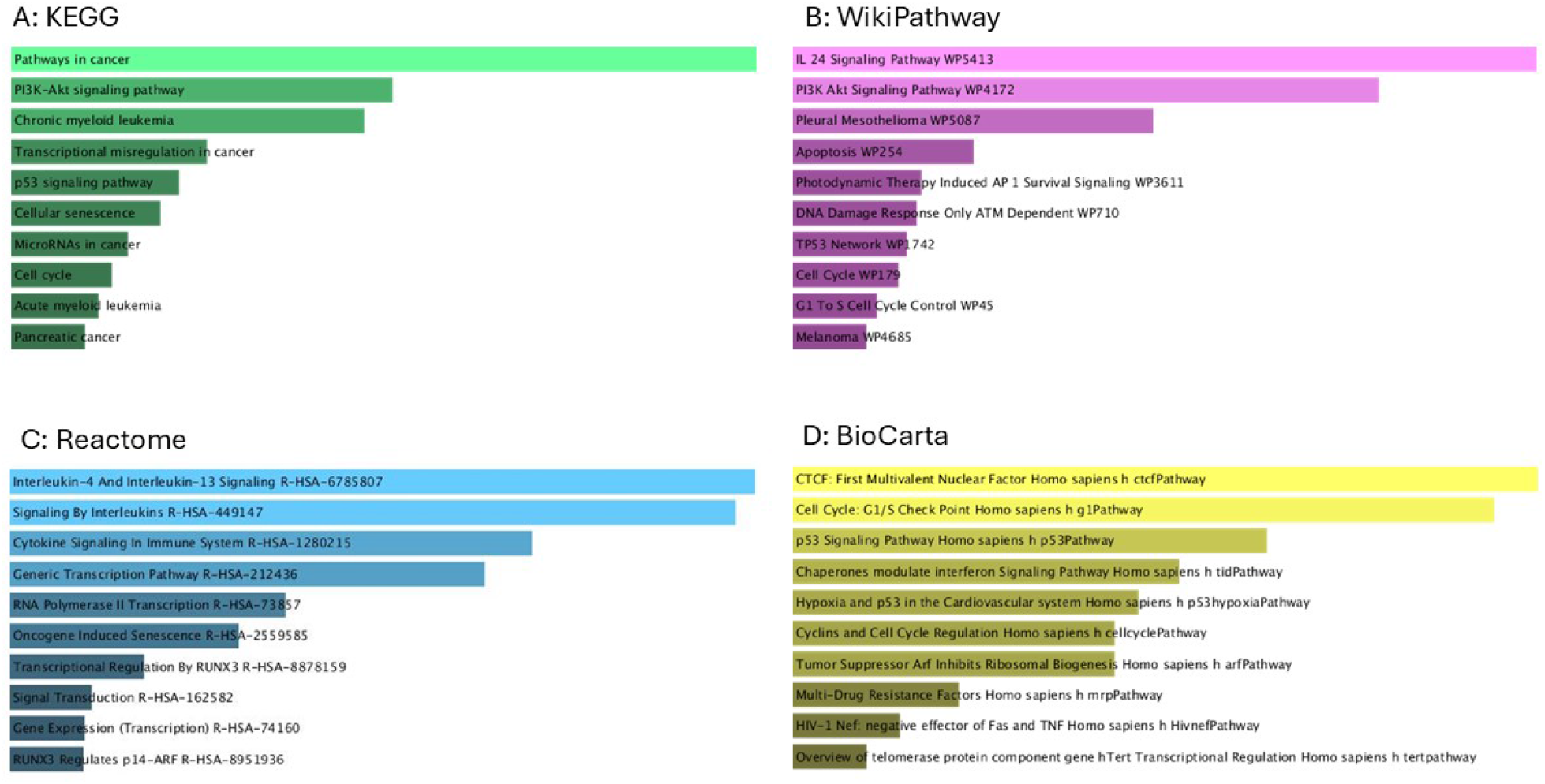
Outcome of pathway enrichment analysis for the common genes using Enrichr programme considering databases: (A) KEGG (B) WikiPathway (C) Reactome and (D) BioCarta.

### Mutation analysis

We performed a mutation analysis on the 15 genes associated with AML and BC CML available on the COSMIC database. This analysis enables us to identify and characterize somatic mutations that may contribute to disease progression and therapy resistance. A detailed information about the known mutations of the common genes, their involvement in different types of tumours, their role in cancer, and their mutation types are summarized in the Table 2 and supplementary table.

**Table 2.**
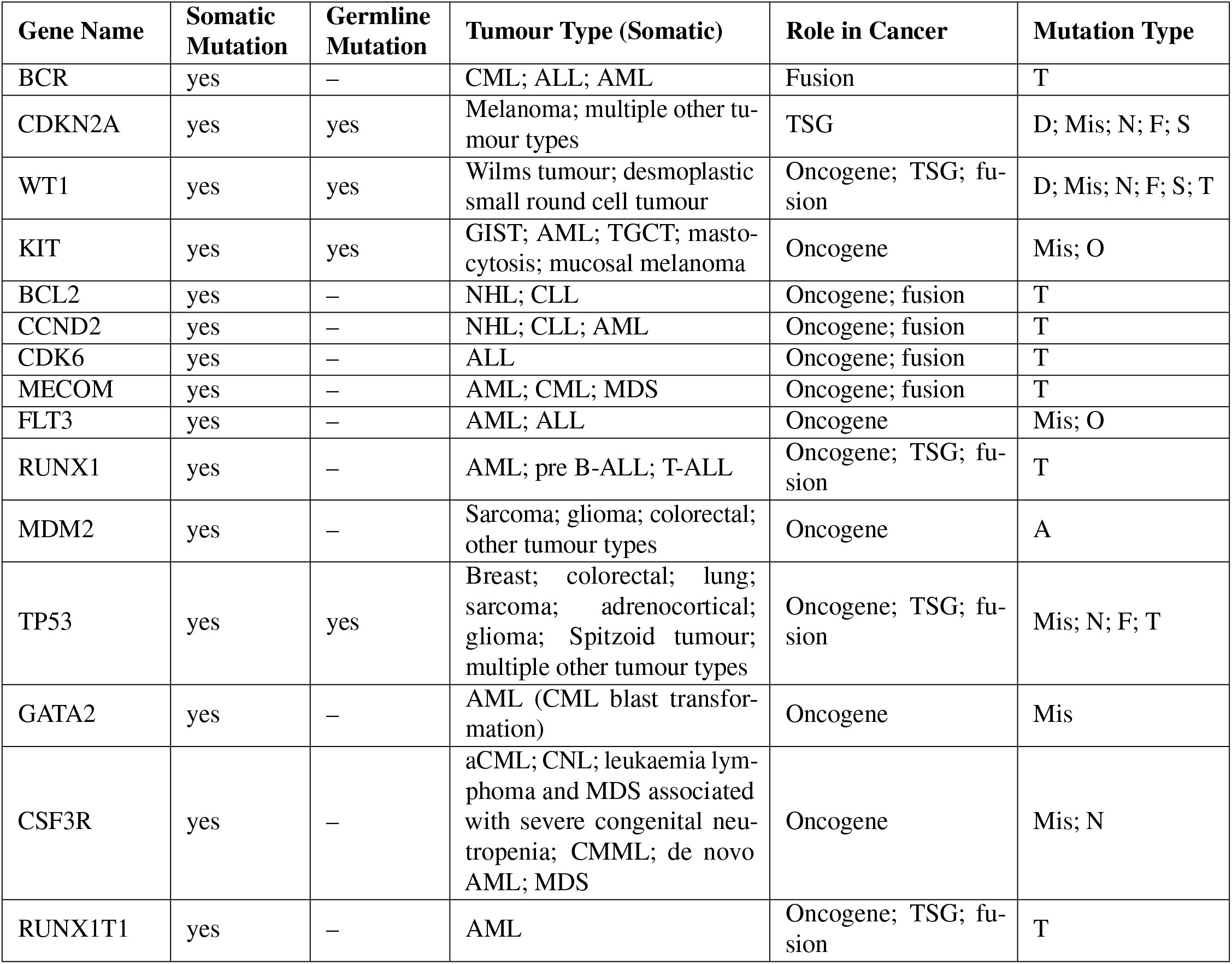
A curated list of mutations, types and their role in cancer for common genes from COSMIC database.

| Gene Name | Somatic Mutation | Germline Mutation | Tumour Type (Somatic) | Role in Cancer | Mutation Type |
| --- | --- | --- | --- | --- | --- |
| BCR | yes | – | CML; ALL; AML | Fusion | T |
| CDKN2A | yes | yes | Melanoma; multiple other tumour types | TSG | D; Mis; N; F; S |
| WT1 | yes | yes | Wilms tumour; desmoplastic small round cell tumour | Oncogene; TSG; fusion | D; Mis; N; F; S; T |
| KIT | yes | yes | GIST; AML; TGCT; mastocytosis; mucosal melanoma | Oncogene | Mis; O |
| BCL2 | yes | – | NHL; CLL | Oncogene; fusion | T |
| CCND2 | yes | – | NHL; CLL; AML | Oncogene; fusion | T |
| CDK6 | yes | – | ALL | Oncogene; fusion | T |
| MECOM | yes | – | AML; CML; MDS | Oncogene; fusion | T |
| FLT3 | yes | – | AML; ALL | Oncogene | Mis; O |
| RUNX1 | yes | – | AML; pre B-ALL; T-ALL | Oncogene; TSG; fusion | T |
| MDM2 | yes | – | Sarcoma; glioma; colorectal; other tumour types | Oncogene | A |
| TP53 | yes | yes | Breast; colorectal; lung; sarcoma; adrenocortical; glioma; Spitzoid tumour; multiple other tumour types | Oncogene; TSG; fusion | Mis; N; F; T |
| GATA2 | yes | – | AML (CML blast transformation) | Oncogene | Mis |
| CSF3R | yes | – | aCML; CNL; leukaemia lymphoma and MDS associated with severe congenital neutropenia; CMML; de novo AML; MDS | Oncogene | Mis; N |
| RUNX1T1 | yes | – | AML | Oncogene; TSG; fusion | T |

### Network construction and interaction between common gene list

The twenty nine common genes were supplied as input in Cystoscope and using STRING database a PPI network with 148 edges of common genes is built and shown in Figure 4. The network file is reintroduced in Cytoscape for visual representation and finding of top hub genes using the CytoHubba plugin of Cystoscope. An analysis of obtained PPI network has identified TP53, ANXA5, RUNX1, IFNG, BCL2, KIT, FLT3, TGFB1, CDKN2A, and MDM2 as the top 10 hub proteins, ranked by their degree of connectivity. However, when ranked by maximum clique centrality, FLT3 is replaced by CCND2 in the top 10 subnetwork (Fig.5). These changes in the subnetwork highlight the differing roles of genes in the network topology and their varying levels of involvement in the overall protein interaction landscape.

**Figure 4.**
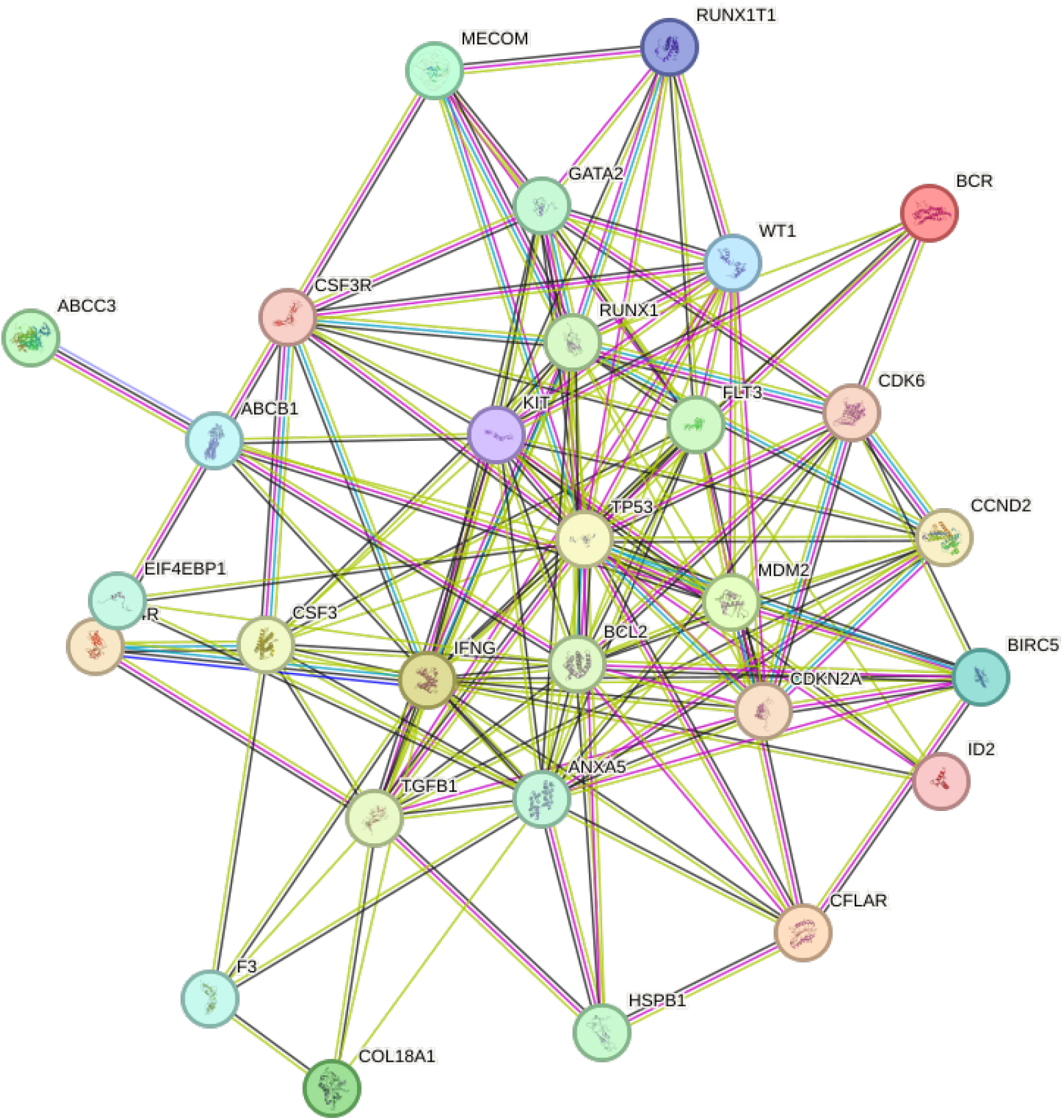
A Protein-Protein Interaction network of common genes among BC CML and AML. The analyzed network holds 29 nodes and 148 edges.

**Figure 5.**
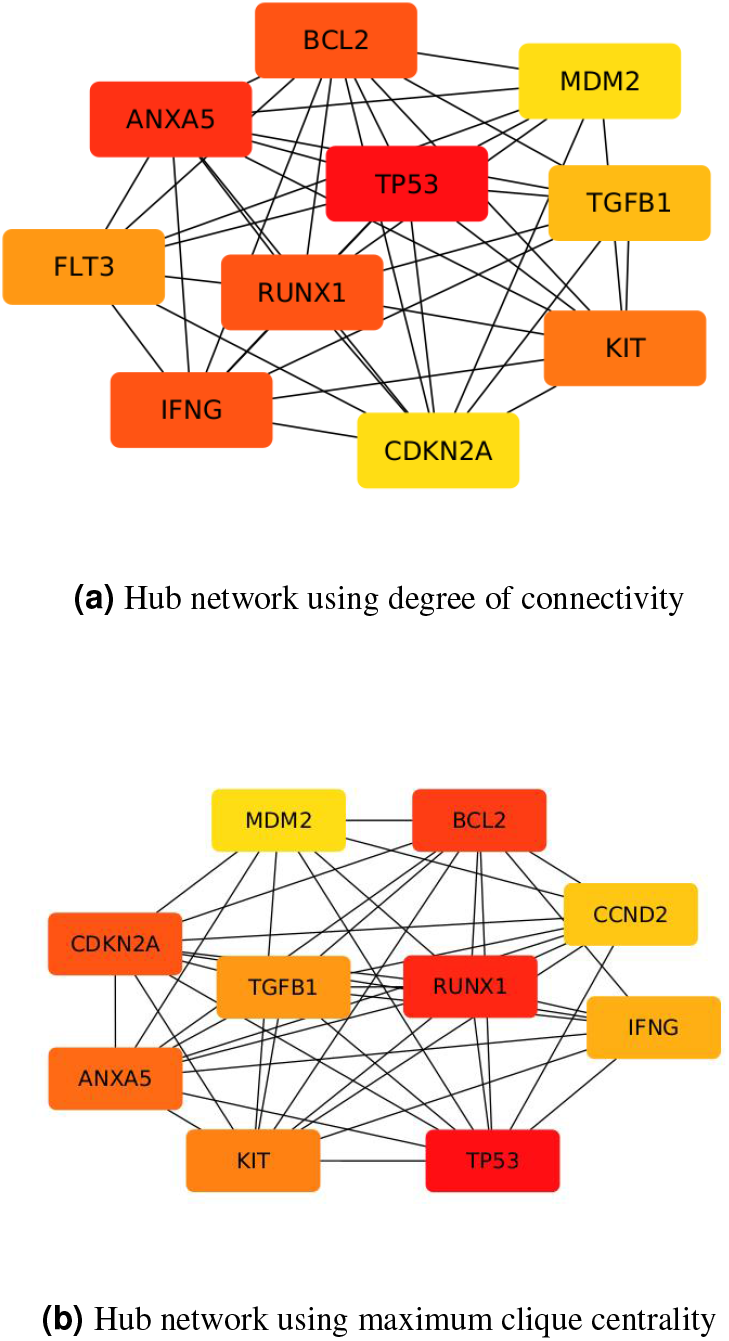
Construction of subnetwork of hub genes for the 29 common genes using cytoHubba

### TF and miRNA mediated network of common genes

A TF-gene interactions is constructed using NetworkAnalyst that displays relationships between 29 common genes and TF genes (Fig.6). We find that VEGFA is the top-ranked transcription factor, followed by TP53 and BCL2. Additionally, these TFs interact with other transcription factors outside of our common gene list, such as SP1, RELA, and NFKB, to regulate downstream pathways in BC CML and AML cells. A miRNA-gene interaction network is also constructed and analysis of that suggests that MDM2 has the highest number of innteractions with miRNAs (degree-205) followed by CDK6 and CCND2. Out of the list of miRNAs, hsa-mir-16-5p had the highest degree of 10 interactions Fig.7. The highly interconnected network suggests significant crosstalk occur among miRNAs in regulating the common genes.

**Figure 6.**
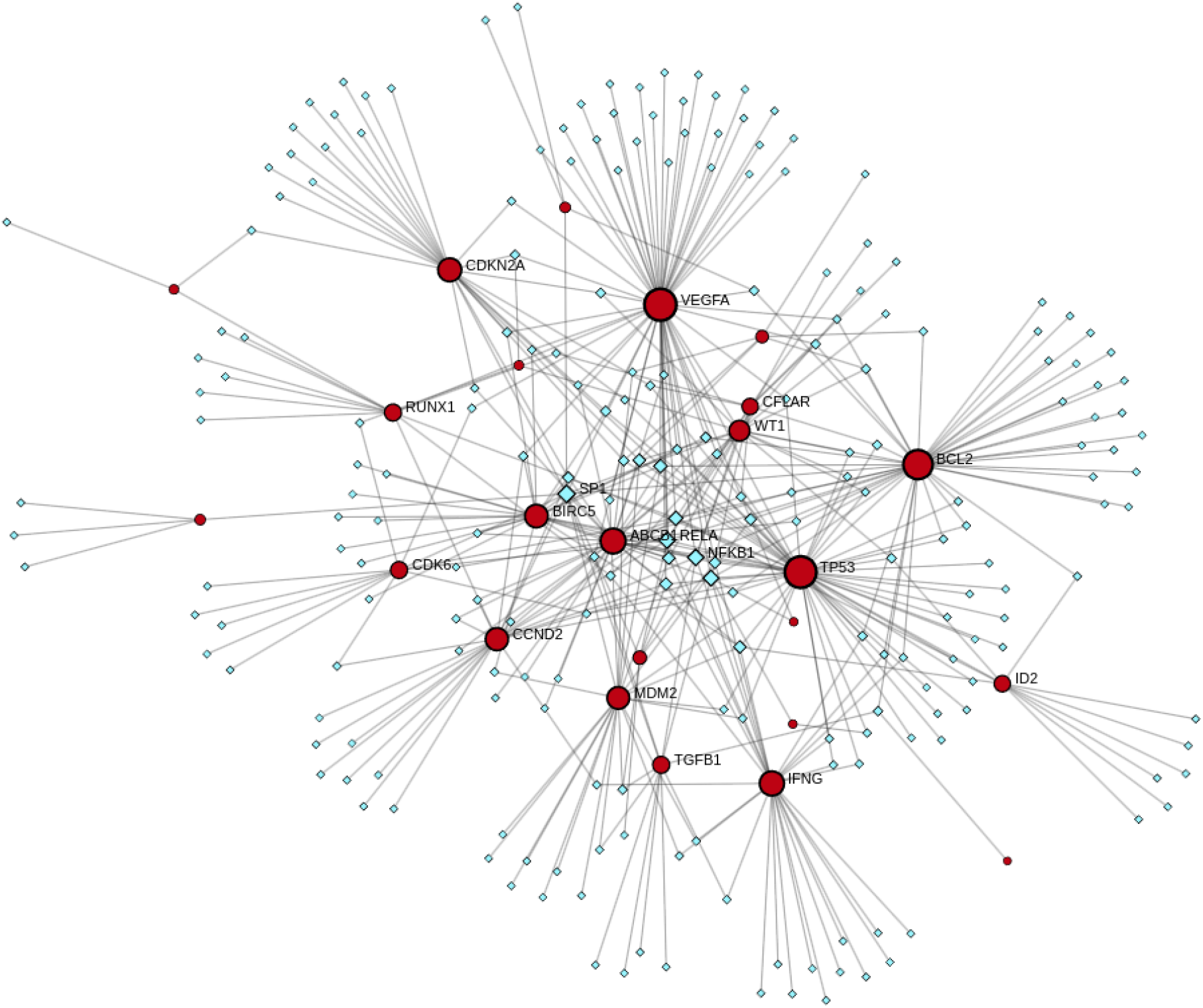
A transcription factor-gene interaction network constructed for all 29 common genes associated with BC CML and AML using NetworkAnalyst.

**Figure 7.**
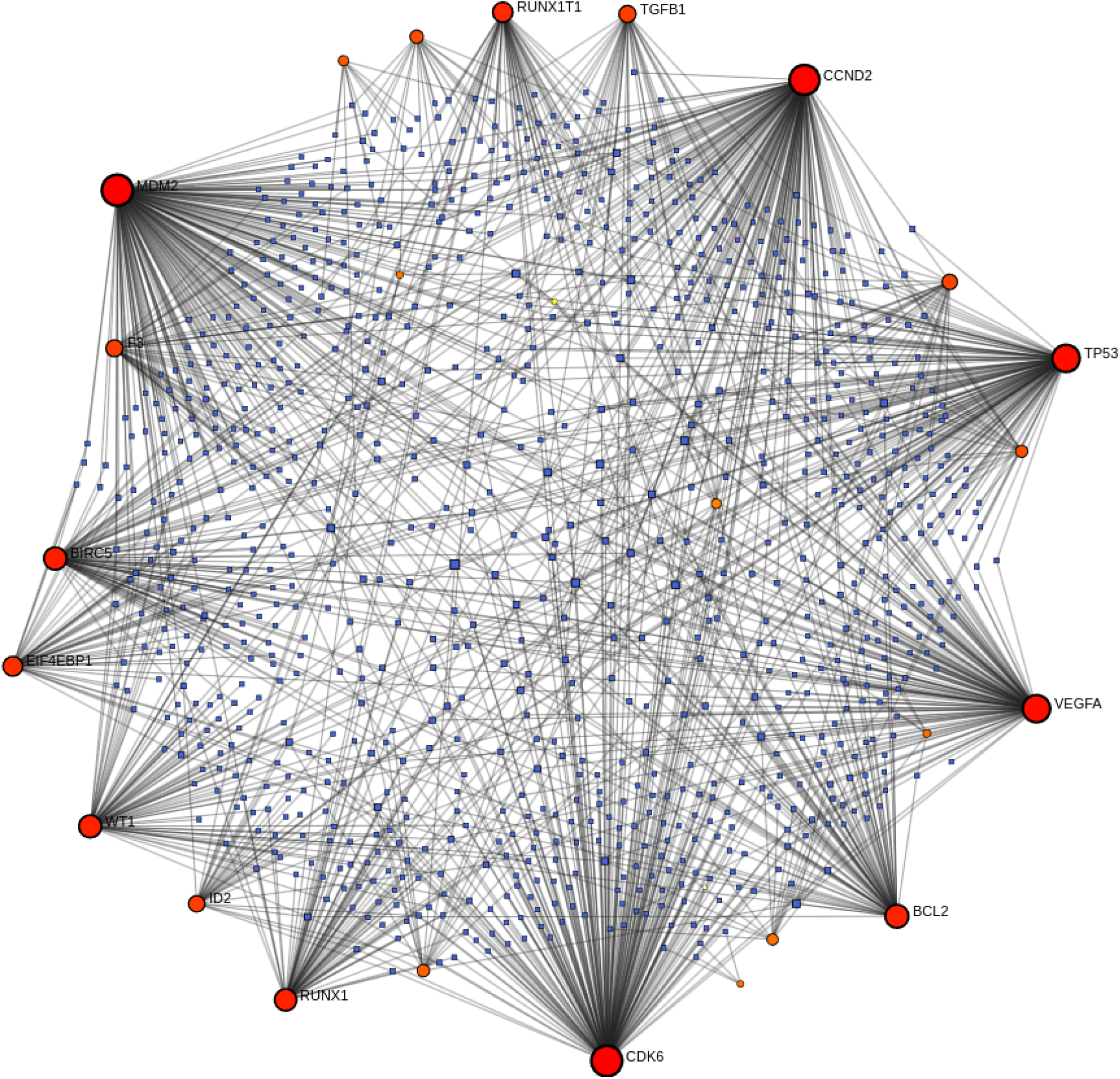
A miRNA-gene interaction network for all 29 common genes associated with BC CML and AML.

### Putative Coregulation in AML and BC CML

Using NetworkAnalyst a TF–miRNA coregulatory network is also generated which is comprise of 240 nodes and 327 edges (Fig. 8). We observe that a total of 130 TF genes and 102 miRNAs are interacting with the 29 common genes.

**Figure 8.**
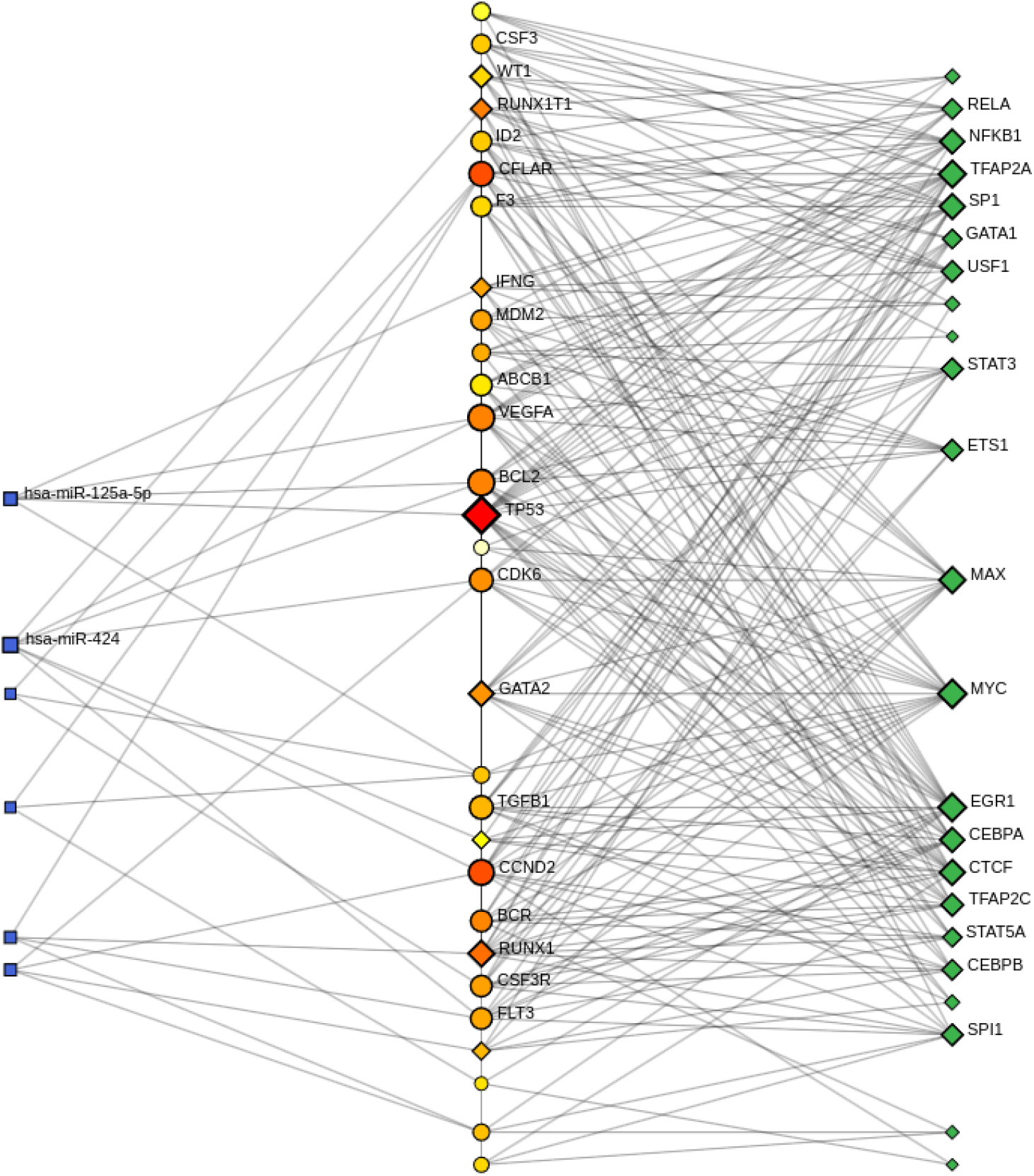
A co-regulatory network depicting the co-regulation of common genes by several Transcription factors and miRNA.

## Discussions and Conclusions

Clinical and pathological similarities between Acute Myeloid Leukemia and Blast Crisis Chronic Myeloid Leukemia have been already explored comprehensively^35^. Both diseases are characterized by a block of differentiation, where hematopoietic stem cells fail to mature into functional blood cells, resulting in an accumulation of immature myeloid blasts in the bone marrow and peripheral blood. Recent studies have also pointed the involvement of similar genetic mutations and pathways in their pathogenesis^15^. To further understand the similarities, our research aims to explore whether AML and BC CML share common regulatory molecules, regulatory networks, or somatic mutations that contribute to their similar clinical features. A deeper understanding of molecular regulations, informed by genetic and expression data, could be beneficial for improving therapy outcomes in BC CML patients.

In the present study, we have collected genes associated with both AML and BC CML from various databases and identified a set of common genes. A gene ontology analysis of these common genes revealed their involvement in biological processes such as the positive regulation of cell population proliferation, negative regulation of programmed cell death, and nucleic acid-templated transcription, which are commonly observed in many cancers^36^. As expected for blood cancers, these genes are also involved in hematopoiesis. Disruption of normal hematopoiesis can lead to myeloid leukemogenesis, particularly driving leukemic transformation, and it is likely that some of the identified genes contribute to these pathological processes. An interesting finding was the positive regulation of serine/threonine kinase activity, which may indicate increased cellular activity, potential upregulation of leukemia-specific kinases, enhanced activity in pathways involving these kinases, or the accumulation of additional passenger mutations^37^. Gene ontology also infers involvement of molecular functions such as DNA-binding Transcription Factor Binding, Cytokine and its receptor Activity, Growth Factor Activity, Protein Tyrosine Kinase Activity. Our results also suggest that most of the common genes are of regulatory nature since their cellular locations are Endocytic Vesicle, Nucleus, Cyclin-Dependent Protein Kinase Holoenzyme Complex, and Serine/Threonine Protein Kinase Complex.

Further, our pathway enrichment analysis suggests existence of molecular connections between AML and BC CML as we have observed several common pathways. The most frequently involved pathways are cell cycle regulation and transcription-related pathways particularly, PI3K-AKT pathway that plays a significant role in cell proliferation and survival.Trials targeting PI3K inhibition may hold therapeutic potential in treating acute forms of leukemia by impairing these key oncogenic processes^38^. Another promising pathway from our result is cytokine signaling, especially from the Interleukin family. If KEGG database is used for the pathway enrichment analysis we get cancer-related pathways, such as the p53 signaling pathway, cellular senescence, and miRNA regulation, as well as pathways specific to CML and AML. WikiPathways further identified the IL-24 signaling pathway as prominently involved among the input genes, alongside pathways like Photodynamic Therapy-Induced AP-1 Survival Signaling and DNA Damage Response (ATM-Dependent). Reactome analysis confirmed enrichment in Interleukin-4 and Interleukin-13 signaling^39^, oncogene-induced senescence, and transcriptional regulation by the RUNX3 transcription factor. Additionally, BioCarta results highlights unique pathways including CTCF as a multivalent nuclear factor, chaperones modulating interferon signaling, and the tumor suppressor ARF inhibiting ribosomal biogenesis.

Somatic mutations in cancer are genetic alterations that occur in non-germline cells, meaning they are not inherited but arise spontaneously in an individual’s cells during their lifetime. They can provide significant insights into Cancer Development and Progression, Tumour Heterogeneity, Biomarkers for Diagnosis and Prognosis, Therapeutic Targets, Resistance Mechanisms, Evolutionary Insights etc. Several genes, such as BCR, MECOM, and GATA2, have known mutations in both AML and CML listed in COSMIC database . Interestingly, GATA2 mutations have been proposed to be specifically associated with blast transformation in CML along with the increase in undifferentiated blast cells observed in both diseases. Mutations in KIT, FLT3, RUNX1, RUNX1T1, and CSF3R genes are predominantly found in AML but not in CML. Mutations in few genes, including BCL2, CCND2, and CDK6, are implicated in other haematological malignancies, such as Chronic Lymphoblastic Leukemia (CLL), Acute lymphoblastic leukemia (ALL), and Myelodysplastic Syndrome (MDS). Additionally, genes like CDKN2A, WT1, MDM2, and TP53 are broadly involved in tumorigenesis across multiple cancer types.

Most identified mutations result from chromosomal translocations in blood cancers, including AML and BC CML. For example, BCR is commonly involved in a translocation with ABL, a well-known driver of CML, but BCR::ABL fusions are also found in some AML patients with poor prognosis^40^. Another notable translocation pair is MECOM::RUNX1. Another main type of mutations found were missense mutations. The roles of these mutations are predominantly oncogenic, with some acting as fusion genes that exhibit both oncogenic and tumour-suppressive functions. We believe that the mutational status of genes which are primarily associated with AML, can be further explored in BC CML. Additionally, mutations in genes involved in other blood cancers may also provide further insights. Comparing mutational signatures between BC CML and AML could provide valuable prognostic biomarkers and inform potential drug responses.

Our protein-protein interaction (PPI) network contained 148 edges, significantly more than the expected number for a network with 29 nodes (43 edges), predicted by the STRING algorithm. This indicates that the proteins in our network interact more frequently with each other than would be expected for a random set of proteins of the same size and degree distribution from the genome, suggesting that these proteins are indeed biologically interconnected.

Using the list of common genes we performed a transcription factor - gene network analysis which indicates that both AML and BC CML are highly dependent on transcription factor regulation. A majority of the 29 common genes are transcription factors, and the the top TFs enriched results from NetworkAnalayst with the highest degree of connectivity in the network are all from our common gene list.

Among the common genes, MDM2 is predicted to be regulated by the largest number of miRNAs, with the highest number of interactions in the miRNA-gene network built by NetworkAnalyst.The MDM2-p53 axis is a well-known miRNA-regulated pathway in oncogenesis, but its role warrants further exploration in the context of AML and BC CML^41^. Also the most connected miRNA is hsa-miR-16-5p which is consistent with the observation that hsa-miR-16-5p has role in carcinogenesis^42^.

TFs and miRNAs play a pivotal role in a variety of biological processes and disorders because they have the ability to both co-regulate the expression of target genes and to mutually regulate one another. To further explore the relation amongst common genes, TFs, and miRNAs, we have constructed a network that reflects co-regulation of our list of common genes by TFs and miRNAs.

There are a number of limitations to this study that need to be recognized and taken into account, such as there aren’t many datasets available for CML, particularly Blast Crisis stage, compared to the wealth of data collected for AML. Also, the genes specifically involved in BC CML are not listed in some databases separately. This could be because of the high success rate of imatinib treatment in chronic phase CML patients leading to a limited population progressing to Blast Crisis. Furthermore, the biological functions and hub gene enrichment analysis results need to be validated by additional experiments and clinical trials since our study is based solely on computational analysis without clinical substantiation.

Finding better alternative therapies is important, especially in the cases with unfavorable TKI response or TKI resistance after certain duration of treatment. Targeted therapies like RUNX1, BCL2, FLT3 inhibitors used in AML against specific mutations can be repurposed in BC CML patients with same mutational profile.

## Acknowledgments

SB acknowledges the Prime Minister Research Fellowship awarded by the Ministry of Education, Government of India for financial assistance. RP acknowledges the Department of Science and Technology, India, for the DST-INSPIRE Faculty Award (DST/INSPIRE/04/2015/001939) and Banaras Hindu University for the Institute of Eminence seed grant. RP also acknowledges the University Grant Commission, India, for start-up grant awarded to him.

## Author contributions statement

SB conceived and performed the analysis. SB and RP analyzed results and prepared the manuscript.

